# BET proteins are required for transcriptional activation of the senescent beta cell secretome in Type 1 Diabetes

**DOI:** 10.1101/736231

**Authors:** Peter J. Thompson, Ajit Shah, Charalampia-Christina Apostolopolou, Anil Bhushan

**Author notes:** These authors contributed equally (Peter J. Thompson, Ajit Shah).

## Abstract

**OBJECTIVE:** Type 1 Diabetes (T1D) results from progressive loss of pancreatic beta cells due to autoimmune destruction. We recently reported that during the natural history of T1D in humans and the female nonobese diabetic (NOD) mouse model, beta cells acquire a senescence-associated secretory phenotype (SASP) that is a major driver of disease onset and progression, but the mechanisms that activate SASP in beta cells were not explored. The objective of this study was to identify transcriptional mechanisms of SASP activation in beta cells.

**METHODS:** We used the female NOD mouse model of spontaneous autoimmune T1D and ex vivo experiments on NOD mouse and human islets to test the hypothesis that Bromodomain Extra-Terminal domain (BET) proteins activate the beta cell SASP transcriptional program.

**RESULTS:** Here we show that beta cell SASP is transcriptionally controlled by BET proteins, including BRD4. Chromatin analysis of key beta cell SASP genes in NOD islets revealed binding of BRD4 at active regulatory regions. BET protein inhibition in NOD islets diminished not only the transcriptional activation and secretion of SASP factors but also the non-cell autonomous activity. BET protein inhibition also decreased the extent of SASP induction in human islets exposed to DNA damage. The BET protein inhibitor iBET-762 prevented diabetes in NOD mice and also attenuated SASP in beta cells in vivo.

**CONCLUSIONS:** Taken together, our findings support a crucial role for BET proteins in the activation of the beta cell SASP transcriptional program. These studies suggest avenues for preventing T1D by transcriptional inhibition of SASP.

## 1. Introduction

Type 1 Diabetes (T1D) is a chronic metabolic disease of insulin deficiency caused by an organ-specific autoimmune disorder that leads to progressive loss of pancreatic beta cells. The conventional model for T1D pathogenesis posits a loss of peripheral tolerance resulting beta cell-specific autoimmunity of CD4^+^ and CD8^+^ T cells that carry out beta cell destruction with support from innate immune cells [1,2]. While T cell-mediated beta cell destruction is indeed the main driver of the disease, we recently demonstrated that beta cells actively participate in the process by undergoing DNA damage-induced senescence, upregulation of the pro-survival factor Bcl-2 and acquiring a senescence-associated secretory phenotype (SASP) [3]. Selective ablation of senescent beta cells with small molecule senolytic compounds targeting Bcl-2 prevented T1D in the NOD mouse model by halting the autoimmune destruction process and preserving beta cell mass [3]. These results demonstrate a causal role for SASP beta cells in the pathogenesis of T1D, however the mechanisms of SASP activation in beta cells have not been determined.

SASP is controlled at multiple levels, including the level of chromatin and transcription [4,5]. Genes encoding SASP factors are dramatically upregulated in senescent cells, which generally correlates with increased protein level and secretion [6,7]. SASP occurs in the context of persistent DNA damage response (DDR) signaling and results from extensive chromatin remodelling and activation of regulatory regions of the genes that encode SASP factors [8–11]. Recent efforts have revealed a growing list of transcription factors, chromatin architectural proteins and modifiers that collaborate to orchestrate the activation of SASP genes, leading to the development of the secretory phenotype. These include factors such as ATM, macroH2A1, HMGB2, MLL1, BRD4, NF-kB, CEBPβ, Mitf, and PARP-1 and inhibition or depletion of these factors in senescent cells reduces SASP gene activation and subsequent protein secretion [5,8–10,12,13]. Activation of SASP genes is also accompanied by changes in chromatin modifications, including the acquisition of marks associated with active enhancers, such as histone H3K27 acetylation (H3K27ac) [8]. Since most of the previous studies investigated the chromatin remodeling and activation of SASP genes in human primary fibroblast lines, it remains unclear whether the some of the same transcription factors and chromatin modifiers are required to activate SASP in other cell types and in vivo. Due to the ability of accumulated SASP cells to profoundly alter tissue microenvironments and disrupt normal processes [14,15], pharmacological approaches to inhibit SASP at the transcriptional level could prove useful for mitigating the effects of senescent beta cells in T1D.

Here we show that transcriptional activation of the SASP in beta cells relies upon epigenetic readers of histone lysine acetylation the BET proteins, including BRD4. BRD4 is expressed in beta cells and was bound to active SASP gene enhancers marked with H3K27ac in NOD islets. Inhibition of BRD4 chromatin binding with the potent small molecule BET inhibitor iBET-762 led to diminished SASP gene expression, protein secretion and SASP paracrine activities from NOD islets. We further show that BRD4 is also expressed in human beta cells and that BET protein transcriptional activity is required for SASP in human islets undergoing DNA damage. Finally, treatment of NOD mice with iBET-762 prevented diabetes and attenuated SASP in beta cells in vivo. Taken together, these results demonstrate that BET proteins are key transcriptional activators of SASP in beta cells and indicate that the mechanism of diabetes prevention with BET inhibitors involves attenuation of SASP.

## 2. Materials and Methods

### 2.1 Animal husbandry and procedures

All procedures involving mice were performed according to the IACUC standards following ethics approval by the animal committee at the University of California San Francisco. Female NOD/ShiLtJ mice (from Jackson Labs) or NOD/MrkTac (from Taconic Labs) were purchased at 8 weeks of age and allowed to age further in our colony. Mice were housed under standard conditions with a 12 hour light-dark cycle. To measure glycemia, 1 mm of tail was clipped and blood was tested with a glucometer and disposable test strips (Freestyle Lite). Mice were considered hyperglycemic with a single blood glucose measurement of >150 mg/dl for fasting blood glucose (FBG) or two consecutive measurements on different days of >300 mg/dl for random blood glucose (RBG). For the diabetes prevention study, euglycemic NOD/MrkTac mice (n = 20) at 12 weeks of age were split into two groups of 10 where one group was injected i.p. with iBET-762 (30 mg/kg) dissolved in DMSO as stock solution of 50 mg/ml, which was further diluted in saline to a concentration of 5 mg/ml. For the control group, an equivalent volume of vehicle (10% DMSO in saline) solution was injected. For iBET and vehicle groups, the total volume injected was between 150-200 µl. The treatment was carried out for 14 consecutive days. Following this, both groups of mice were monitored for RBG. The RBG was measured once per week until they reached 30 weeks of age. At the end of the study, pancreata from the remaining euglycemic mice were harvested, formaldehyde-fixed and paraffin-embedded. For assessment of insulitis, 5 µm paraffin-embedded sections from two vehicle and two iBET762-injected mice at 30 weeks of age from regions 50 µm apart in the pancreas block were stained with hematoxylin and eosin. Insulitis was scored for three different stages based on the level of infiltration of islet by immune cells, no insulitis, peri-insulitis (infiltration at the periphery of the islets), and insulitis (infiltration within the islet) with n = 30 islets were scored per mouse. For the iBET treatment of hyperglycemic mice, once the NOD/MrkTac were hyperglycemic as defined by RBG > 300 mg/dl they were injected i.p. with iBET-762 (30 mg/kg) or vehicle, for seven consecutive days then sacrificed for pancreas dissection and histology.

### 2.2 Human pancreas sections and islets

Human pancreas sections were obtained from the Network for Pancreatic Organ donors with Diabetes (nPOD) program [16]. nPOD pancreas donors used in this study were nPOD Case IDs: 6279 (nondiabetic, 19-year old male), 6397 (GADA autoantibody-positive 21-year old female), 6342 (IA-2A, mIAA autoantibody-positive, T1D 14-year old female). Human islets were obtained from the Integrated Islet Distribution Program (IIDP). Human islet preparations used in this study were: Donor #1: RRID:SAMN10784981, 45 year old female, BMI = 35.7 and; Donor #2: RRID:SAMN11046361, 57 year old male, BMI = 35.9.

### 2.3 Culture of mouse islets, inhibitors and RNAi

Islets from NOD/ShiLtJ mice were isolated according to standard methods by collagenase digestion through the bile duct. For collection of conditioned media (CM) from whole islets, after 1-2 h of resting at 37°C, NOD islets from n = 3 mice were separated into two equal pools per mouse, where one half were cultured in with vehicle (0.1% DMSO) and the other half were cultured in 5 µM iBET-762 (ApexBio) in non-treated 96-well plates at 37°C with 5% CO_2_ in 200 µl of serum-free mouse islet media (RPMI 1640, 2 mM L-glutamine, 1X penicillin-streptomycin) for 24 h. Following the incubation, CM was collected for analysis and islet DNA was either extracted immediately (Zymogen total genomic DNA kit) or islets were stored at −80°C in genomic DNA lysis buffer to be extracted later. Collection of CM from CD45-depleted islet cell cultures was performed as previously [3] where islets were pooled from five or ten 14 week NOD mice, depleted of CD45^+^ immune cells using MACS (BioLegend) and plated at 60,000-100,000 cells per well (equal numbers per well) in 3 biological replicates per group in mouse serum-free islet media containing vehicle (0.1% DMSO) or 5 µM iBET-762 for 24 h. Following incubation the CM was collected for paracrine assays. For qRT-PCR of CD45-depleted islet cultures, cells were incubated with iBET-762 in media containing 10% FBS (rather than in serum-free media). For RNAi, after recovery of NOD islets for 18 h in serum-containing islet media (RPMI 1640, 2 mM L-glutamine, 1X penicillin-streptomycin, 10% FBS) 20 islets per well (n = 3 biological replicates per group) were transfected with control (non-targeting #2) siRNA or mouse *Brd4* SMARTpool siRNA (Dharmacon GE Life Sciences) at 100 nM each with lipofectamine 3000 according to the product instructions. Islets were cultured an additional 72 h and then harvested for RNA extraction. THP-1 human monocytic leukemia cells (purchased from ATCC) were cultured in RPMI-1640, 10% FBS and 1X pennicillin-streptomycin at 37°C with 5% CO_2_.

### 2.4 Culture of human islets and inhibitors

Human islets from IIDP were cultured as previously [3] in RPMI-1640 containing 10% FBS, 5.5 mM glucose and 1X antibiotic-antimycotic (Gibco). Islets were rested for 24 h in culture prior to inducing DNA damage with 50 µM bleomycin. Forty-eight hours later bleomycin was removed and fresh media was added, and islets were then cultured with vehicle (0.1% DMSO) or 5 µM iBET-762. Five days later islets were transferred into serum-free media for the collection of conditioned media (CM). After 24 h, CM was collected for luminex assays and islets were harvested for RNA extraction and qRT-PCR.

### 2.5 Immunohistochemistry and quantification

Immunohistochemistry of formaldehyde-fixed paraffin-embedded pancreas sections was performed as described [17]. Five micron tissue sections were rehydrated with xylene and graded ethanol washes, soaked in 1% H_2_O_2_ for 10 minutes, and subjected to heat-mediated antigen retrieval with sodium citrate pH 6.0 for 7.5 minutes at 100% and 14 minutes at 50% power in a 1250W microwave. Sections were cooled to room temperature with tap water and then permeabilized with TBS containing 0.1% Triton-x-100 for 5 minutes. Sections were blocked with 2% normal donkey serum in protein block buffer (Dako) for 15 minutes and then incubated overnight at 4°C with primary antibodies diluted in DAKO antibody diluent: anti-Insulin (DAKO, 1:1000), anti-BRD4 (Bethyl laboratories, 1:10,000), anti-IL-6 (Novus Biologicals 1:300), anti-Flnb (Novus Biologicals, 1:500) [3]. FITC or Cy3-conjugated secondary antibodies (Jackson Immunoresearch) were used to detect primary antibodies and sections were counterstained with DAPI (Vectashield). Images were taken on a Zeiss Axioscope2 wide-field fluorescence microscope with Axiovision software. IL-6 was quantified in beta cells from recent onset diabetic NOD mice ranging in ages from 12-19 weeks that were vehicle treated (n=3) or iBET-762 treated (n=3), with three pairs of age-matched mice among the groups (12-14 week, 16 week, 17-19 week). Since the mice were diabetic in this study, they had widely varying numbers of remaining islets and remaining beta cells per islet. To account for these differences, frequencies of IL-6^+^ beta cells were calculated from a similar number of total beta cells scored from the mice. A total of n = 20 islets with 735 beta cells were scored from vehicle mice, and n = 16 islets with 891 beta cells were scored in iBET-762 mice taking sections throughout the pancreas from mice in both groups. Residual islets containing fewer than 10 beta cells were not scored.

### 2.6 THP-1 chemotaxis assay

Chemotaxis assays were performed as previously described using transwell inserts with 5 µm polycarbonate membranes [3]. Briefly, 1×10^5^ THP-1 cells were seeded into 150 µl of control media (serum-free THP1 media: RPMI 1640, 2 mM L-glutamine, 1% penicillin-streptomycin) in the upper chamber and 150 µl of control media alone or islet CM (with vehicle or iBET-762 treatment) was added to the lower chamber. Cells were allowed to migrate for 5-6 hours at 37°C and then the inserts were removed and the cells in the lower chamber were counted. The control media in the lower chamber was used to determine background levels of migration and the fold-change relative to the control media was used to determine relative chemotaxis in the other samples.

### 2.7 Paracrine senescence assays

Paracrine senescence assays were performed as previously described [3]. NOD islets isolated from 3-week old mice were cultured for 4 days with CM from 14-15 week CD45-depleted islet cell cultures treated with vehicle or iBET762 and then harvested for RNA and qRT-PCR.

### 2.8 ChIP and qPCR analysis

Crosslinked ChIP was performed with the MicroChIP kit (Diagenode) according to the product instructions. For H3K27ac ChIP, the islets were prepared from 13-14 week old NOD/MrkTac or NOD/ShiltJ mice and rested 18 h in mouse islet media. The following day, all the islets from one mouse each were dissociated into single cell suspension using a nonenzymatic buffer reagent (Gibco) and counted with trypan blue. Approximately 2-5×10^5^ cells were crosslinked with 1% formaldehyde for 10 minutes at room temperature, quenched with glycine, then sonicated with the Diagenode Bioruptor to generate ∼200-600 bp fragments. ChIP was performed with 1 µg of anti-H3K27ac (Abcam ab4729) or total rabbit IgG (Sigma-Aldrich) and incubating overnight at 4°C, followed by capture on protein A dynabeads. Crosslinked ChIP of BRD4 from NOD mouse islets was performed using the True MicroChIP kit (Diagenode) and the same procedure was followed for preparing islets. Then instead of resting overnight, the islets from five NOD mice were divided into two pools, where one set were incubated with vehicle (0.1% DMSO) and the other incubated with 5 µM iBET-762 for 18 h (n = 2 biological replicates per group). Islets where then dissociated, crosslinked, and sonicated as described above. ChIP was performed using 1 µg of anti-BRD4 (Bethyl labs A301-985) or total rabbit IgG (Sigma-Aldrich). Ten percent input fractions from all were saved and DNA extracted in parallel with the ChIP samples for qPCR normalization. qPCR using SYBR green was performed using the following forward and reverse primers (5’-3’): *Mmp2* enhancer AACATGCAAGGGAGATCAGC and 5’GGGCTATGAAATGCCAGAAA; *Mmp2* distal GCTGGTCCCACAGTGAAGTT and GGACACAGCGCAACTGAATA; *Il6* promoter TGTGGGATTTTCCCATGA and TGCCTTCACTTACTTGCAGAGA; *Il6* distal GAGAGAGGCAAGCACAGAAA and CCGAGCTGGTTGGTGATAAG. ChIP enrichment was calculated using the percent input method.

### 2.9 Quantitative Reverse-transcriptase PCR

Total RNA was extracted from mouse or human islets cultured as described above and treated with DNAse I with the Zymogen RNA extraction micro kit (Zymogen) according to the product instructions. RNA was converted into cDNA using oligo(dT) primer and the SuperScript IV kit (Invitrogen) or the qScript kit (QuantaBio). qPCR was performed with the ABI 7900HT system and SDS 2.4 software and target gene primers used previously, and relative quantification was performed with the ΔΔC_T_ method using RQ manager 1.2.1, and expression values were normalized to mouse *Ppia* or human *PPIA* as a housekeeping gene [3]. Primer sequences for mouse *Il6, Ppia*, and human *CDKN1A, CDKN2A* (p16), and *PPIA* were as previously [3]. Newly designed forward and reverse primer sequences used in this study were (forward and reverse, 5’-3’): *Flnb*: AACCAGAACTGGAAGATGG and ATGGCTTCTCGGGCATTATC; *Mmp2*: CCCCGATCTACACCTACACC and GGAGTGACAGGTCCCAGTGT; *Igfbp4*: ATCGAAGCCATCCAGGAAAG and CAGGGGTTGAAGCTGTTGTT; and *Brd4*: TTCAGCACCTCACTTCGACC and CTGGTGTTTTTGGCTCCTGC.

### 2.10 Luminex assays

Luminex assays were performed on 50 µl of islet CM, according to the product instructions using either the custom human panel (R&D systems) reported previously [3] or using a custom mouse panel (R&D systems) including: IL-6, Mmp12, Mmp3, Igfbp3. Analyte concentrations were determined from standard curves of purified proteins in each assay provided in the kits. For mouse islet secretion assays involving inhibitor treatments, islets were saved for DNA extraction to normalize concentrations to DNA content and data were pg of secreted factor per µg DNA and expressed as a fold-change relative to the pool of vehicle-treated islets from the same mouse (n = 3 mice). For human islet secretion assays, data were normalized to RNA content and reported as pg secreted factor per µg RNA.

### 2.11 Statistics

The number of biological replicates, number of mice used, or repeated experiments are indicated in the Figure legends. Statistical analyses were performed where experiments were carried out using three or more biological replicates or independent experiments using unpaired one-way ANOVAs for groups of three or with unpaired two-tailed T-tests for groups of two. Log-rank test was used to compare diabetes incidence curves. Results were considered significant at p < 0.05. Statistics were computed using GraphPad prism v6.0.

## 3. Results

### 3.1 BRD4 is required for SASP gene activation in NOD islets

We previously demonstrated the expression of SASP genes in beta cells at 14-16 weeks in euglycemic NOD mice [3]. To confirm that SASP gene regulatory regions were activated in NOD islets, we performed chromatin immunoprecipitation (ChIP) for H3K27ac. Female NOD mice were used throughout our studies due to their higher disease penetrance [18]. We used the dbSUPER superenhancer database [19] and the VISTA enhancer database (Lawrence Berkeley National Lab) to identify putative enhancers of SASP genes from different mouse cell types. ChIP analysis revealed enrichment of H3K27ac within the enhancer of *Mmp2* and the promoter/enhancer of *Il6* relative to distal regions in the islets of some euglycemic NOD mice but not others (Fig. 1a), consistent with the heterogeneity in the extent of beta cell SASP among mice at the same age [3]. Epigenetic readers of H3K27ac include the BET proteins BRD2, BRD3 and BRD4, which are expressed in most somatic cell types and are recruited to regulatory regions to activate transcription. BRD4 is a crucial activator of SASP gene enhancers in human fibroblasts [8,20] and therefore we focused on this paralogue. Immunohistochemistry (IHC) demonstrated the expression of BRD4 in beta cells of euglycemic NOD mice (Fig. 1b). Consistent with the activation state of SASP genes, ChIP analysis revealed that BRD4 was enriched at the same regulatory regions of *Mmp2* and *Il6* that harbored H3K27ac and treatment of islets with the potent small molecule BET inhibitor iBET-762 disrupted its binding at these sites (Fig. 1c). To determine whether BRD4 activates SASP genes in specifically in beta cells, dispersed islets isolated from euglycemic 14-week NOD mice were depleted of CD45^+^ immune cells to enrich for beta cells and then incubated with iBET-762 (Fig. 1d). iBET-762-treated islet cell cultures showed reduced expression of specific beta cell SASP genes *Mmp2, Il6, Flnb* and *Igfbp4* [3] (Fig. 1d). Since iBET-762 can affect the chromatin binding of all three BET paralogues, we also used siRNA knockdown to assess the specific contribution of BRD4. siRNA knockdown of *Brd4* in NOD islets diminished *Il6* expression (Fig. 1e). Together these results implicate BET proteins, and in particular, BRD4 as activators of SASP gene expression in beta cells.

**Figure 1.**
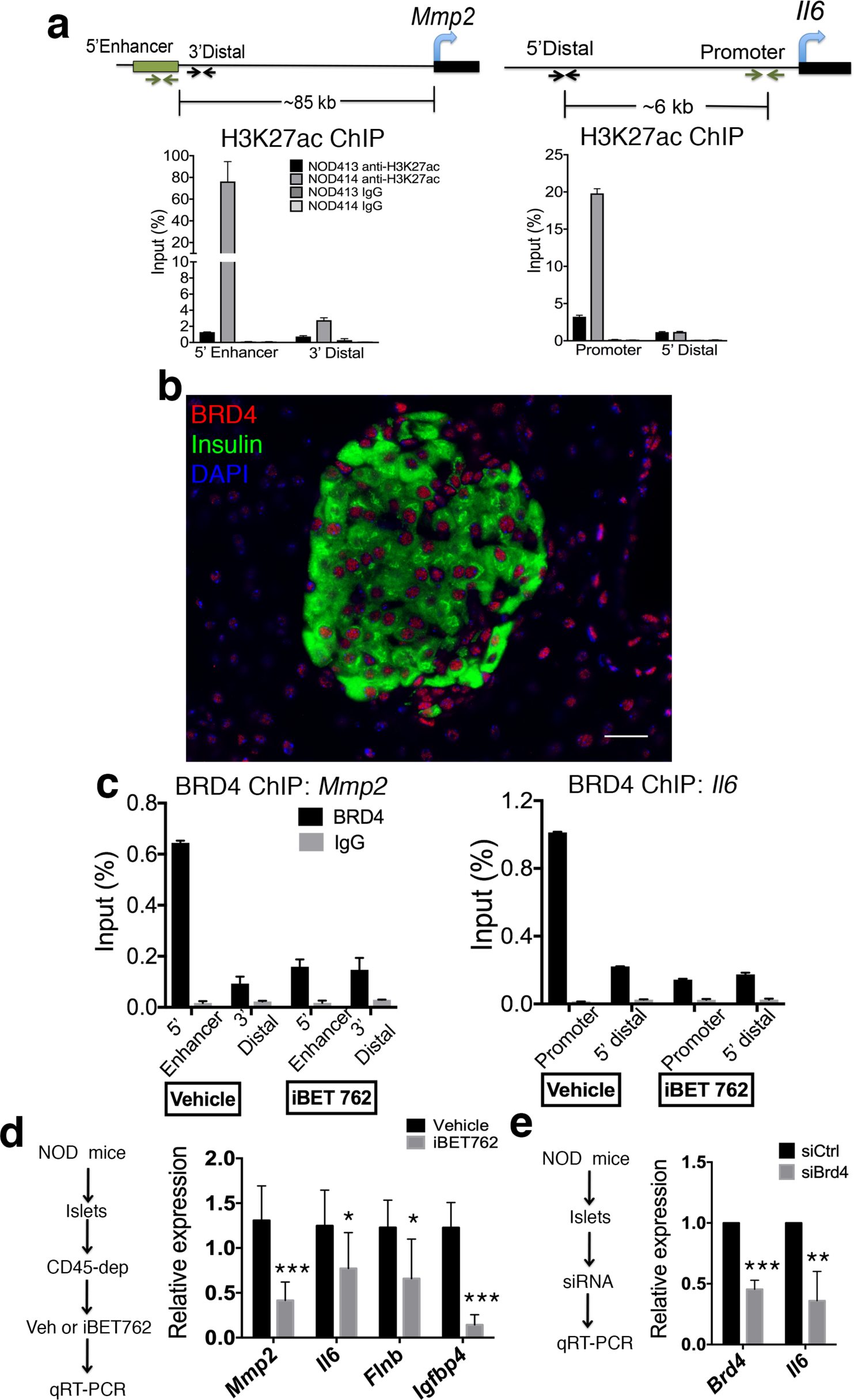
BET protein BRD4 is required for transcriptional activation of SASP genes in NOD islets. (a) ChIP and qPCR for H3K27ac at the enhancer of *Mmp2* and promoter of *Il6*, along with distal negative regions, on islets isolated from two different euglycemic 14-week NOD mice (NOD413, NOD414), n = 2 biological replicates per mouse. Female mice are used throughout our study. The schematic shows the positions of the regions in the ChIP. IgG ChIP was a negative control. Error bars are SD. (b) Representative IHC for BRD4 and Insulin with DAPI on pancreas section from euglycemic NOD mice at 8 weeks of age. Scale bar = 20 µm. (c) ChIP and qPCR for BRD4 at the same regulatory regions of *Mmp2* and *Il6* shown in (a) on islets from euglycemic 14-week NOD mice cultured for 18 h with vehicle or iBET-762 (n= 2 biological replicates per group), error bars are SD. IgG ChIP was a negative control. (d) Islets isolated from 14-week NOD mice were depleted of CD45^+^ to enrich for beta cells and the cultured for 24 h with vehicle (DMSO) or iBET-762 followed by qRT-PCR analysis. qRT-PCR of SASP genes *Mmp2, Il6, Flnb* and *Igfbp4* show the average relative expression for vehicle (n = 5 biological replicates) or iBET-762 treated (n = 4 biological replicates) islets cells. Error bars are SD. (e) qRT-PCR for *Brd4* and *Il6* on 14-week NOD islets transfected with control or *Brd4* siRNAs. Data show average expression levels in siBrd4 relative to siCtrl from n = 3 independent experiments. Error bars are SD. For all panels, *p < 0.05, **p < 0.005, ***p<0.0005, two-tailed T-tests.

### 3.2 BET inhibition blunts SASP secretion and paracrine activities

Next we determined whether disrupting BET protein chromatin binding affected SASP protein secretion. To this end, we cultured NOD islets in iBET-762 or vehicle and collected CM for analysis of SASP factors by luminex assay (Fig. 2a). Notably, iBET-762-treated islets showed significantly lower secretion of SASP factors IL-6, Igfbp3, Mmp3 and Mmp12 (Fig. 2a). Beta cell SASP exerts non-cell autonomous effects including the ability to induce senescence-related changes in immature islet cells and stimulate the chemotaxis of monocytes in vitro [3]. To address whether iBET-762 treatment affected the paracrine senescence activities of SASP, we cultured islets from 3 week NOD mice with CM from islets depleted of CD45^+^ cells of vehicle or iBET-762-treated 14 week NOD mice (Fig. 2b). While CM from vehicle-treated islets showed induction of *Cdkn1a* expression, consistent with paracrine senescence, the iBET-762-treated islet CM did not (Fig. 2b). Finally, we tested whether the CM from iBET-treated NOD islets had effects on monocyte chemotaxis using the transwell assay (Boyden chamber assay) on the human THP-1 monocyte cell line. Importantly, CM from NOD islet cells cultured with iBET-762 showed diminished chemotactic activity relative to the vehicle-treated islet CM (Fig. 2c) in line with their lower SASP factor secretion (Fig. 2a). These data demonstrate that disruption of BET protein transcriptional activity in islets leads to decreased SASP secretion and non-cell autonomous activities.

**Figure 2.**
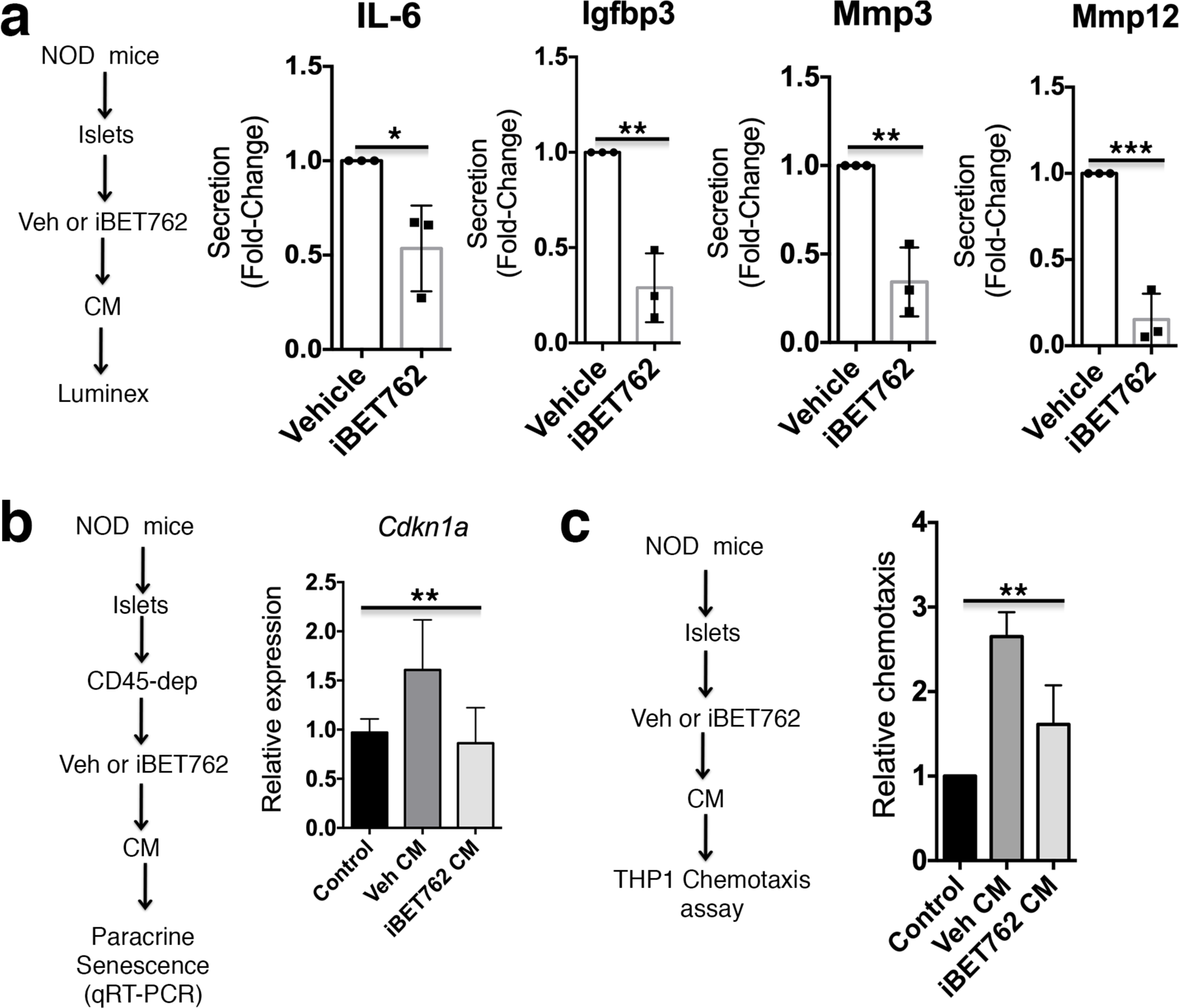
iBET-762 attenuates SASP secretion and paracrine activities from NOD islets. (a) Islets isolated from 14-week NOD mice were cultured in vehicle or iBET-762 for 24h and the conditioned media (CM) was collected for luminex assays. Relative quantification of average secretion of SASP factors IL-6, Igfbp3, Mmp3 and Mmp12 from iBET-762-treated islets versus vehicle-control islets, n = 3 biological replicates per group. Error bars are SD. *p<0.05, **p<0.005, ***p<0.0005, two-tailed T-tests. (b) Islets isolated from 14 week NOD mice were depleted of CD45^+^ cells, and cultured in vehicle or iBET-762 for 24 h and the resulting CM was collected used in paracrine senescence induction assays on islets from 3-week old NOD mice. qRT-PCR for *Cdkn1a* as a readout of paracrine senescence induction in islets from 3-week NOD mice. Regular islet media (not conditioned by islets) was used as a negative control to set the baseline. Data are average relative expression levels from n = 5 or 6 biological replicates per group, error bars are SD. **p < 0.005, one-way ANOVAs (c) CM was collected from islets of 14-week NOD mice cultured with vehicle or iBET-762 and used in chemotaxis assays on THP-1 cells. Regular islet media (not conditioned by islets) was used as a negative control to measure the baseline. Data are chemotaxis fold-change, relative to control, from n = 3 biological replicates per group. Error bars are SD. **p <0.005, one-way ANOVA.

### 3.3 BET proteins are required for SASP in human islets

Induction of DNA damage in human islets ex vivo can lead to acquisition of a SASP reminiscent of the SASP expressed in beta cells of T1D donors [3]. IHC of BRD4 in age-matched human nondiabetic, autoantibody-positive and T1D donor pancreas sections showed that it was expressed in the vast majority of INS^+^ and INS^-^ cells (Fig. 3a). To determine whether BET proteins activate SASP in human islets, we utilized our previous approach for inducing SASP in human islets [3]. Human islets were first treated with bleomycin (bleo) to induce DNA damage or vehicle as a control. Then after washout of bleo or vehicle, islets were cultured in the presence of vehicle or iBET-762 and monitored for persistent DDR and SASP development by qRT-PCR and luminex assays 6 days later (Fig. 3b). Bleomycin treatment induced sustained expression of *CDKN1A* relative to the vehicle control in two different preparations of donor islets (male and female), whereas *CDKN2A* (*INK4A/P16*) was unaffected (Fig. 3c) consistent with previous our findings [3]. In contrast, iBET-762 treatment diminished the expression *CDKN1A* after Bleo (Fig. 3c) suggesting a reduction in the extent of persistent DDR signaling. Furthermore, while Bleo-treated islets developed a SASP as evidenced by upregulated secretion of SASP factors CXCL1, IL-8 and IGFPB4, iBET-762 treatment after bleo attenuated the secretion (Fig. 3d). Taken together these data demonstrate that BET proteins are required for SASP activation in human islets.

**Figure 3.**
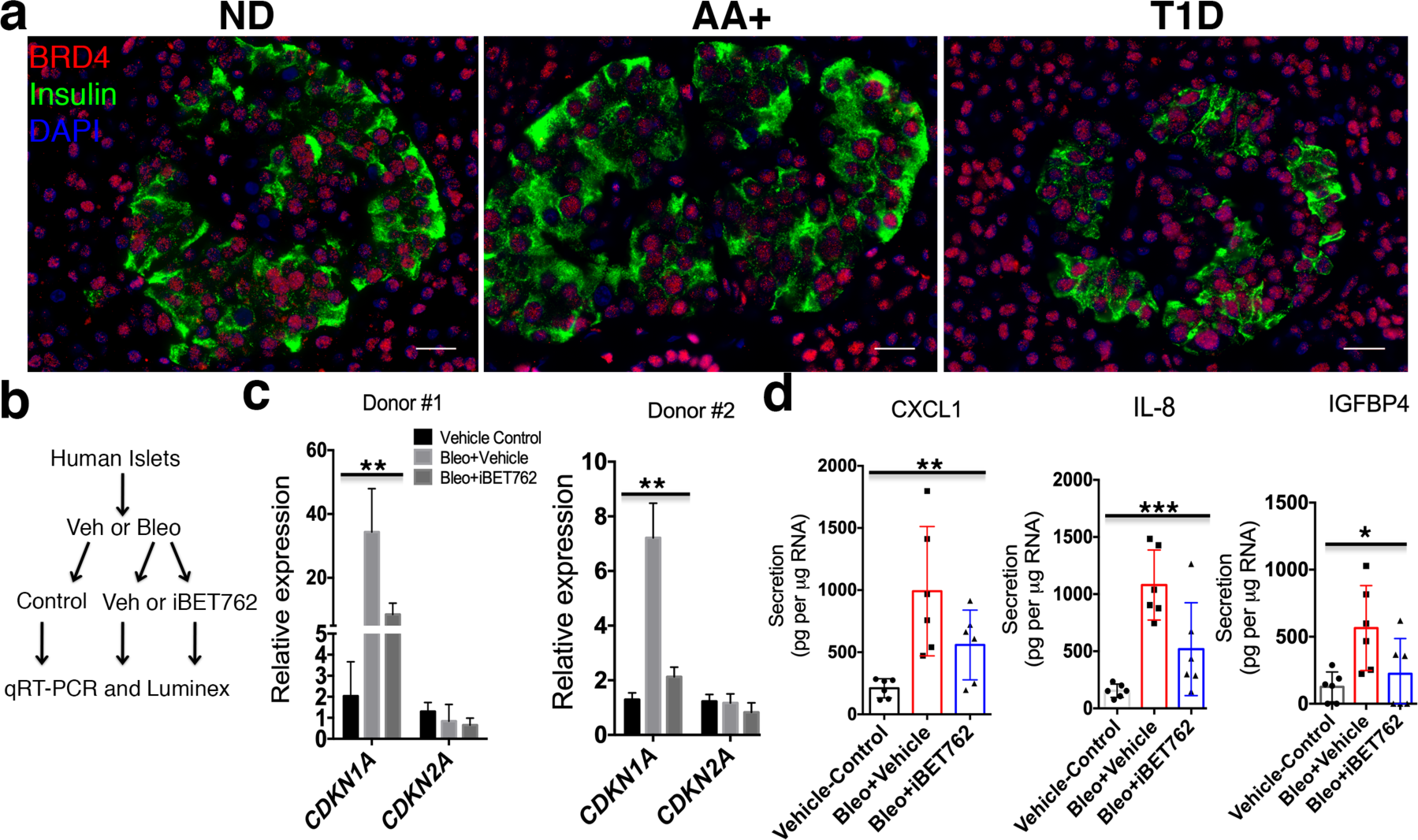
BET proteins are required for activation of SASP in human islets exposed to DNA damage. (a) Representative IHC for BRD4 and Insulin with DAPI in pancreas sections from nondiabetic (ND), autoantibody-positive (AA+) and T1D donors. Scale bars = 20 µm. (b) Islets from human donors were cultured with bleomycin (bleo) to induce DNA damage, or vehicle (0.1% DMSO) as a control, followed by replacing the media with media containing iBET-762 or vehicle and culture for an addition 5 days. Then islets were harested for qRT-PCR and conditioned media (CM) saved for luminex assays. (c) qRT-PCR analysis of *CDKN1A* and *CDKN2A* (*INK4A/P16*) on vehicle control, Bleo+vehicle or Bleo+iBET-762 islets isolated from two different donors (n = 3 biological replicates per group for each donor). Error bars are SD. **p <0.005, one-way ANOVAs. (d) Luminex assay of average protein secretion of SASP factors CXCL1, IL-8 and IGFPB4 from the same islets as in (c). Each dot represents a biological replicate of islets (n = 6 per group, from the two different donors shown in (c). Error bars are SD. *p < 0.05, **p <0.005, ***p<0.0005, one-way ANOVAs.

### 3.4 BET protein inhibition diminishes beta cell SASP in vivo

Previous work has shown that inhibition of BET proteins with the earlier generation inhibitor iBET-151 protects against diabetes in NOD mice [21], however, the mechanisms of its action on beta cells was not determined. To investigate this mechanism, we first treated a cohort of 12-week old euglycemic NOD mice (n = 10 mice per group) with either vehicle or iBET-762 over a two-week period and then monitored random blood glucose levels weekly until 30 weeks of age (Fig. 4a). Blood glucose measurements showed that 60% of vehicle-injected mice developed diabetes by 30 weeks of age whereas only 10% of the iBET-762 injected mice developed hyperglycemia by 30 weeks of age (p = 0.0115, Fig. 4a). Histological staining of pancreas sections confirmed that a greater number of islets were preserved intact in iBET-762-treated mice (Fig. 4b) indicating that iBET-762 halted disease progression, consistent with the previous study using iBET-151 [21].

**Figure 4.**
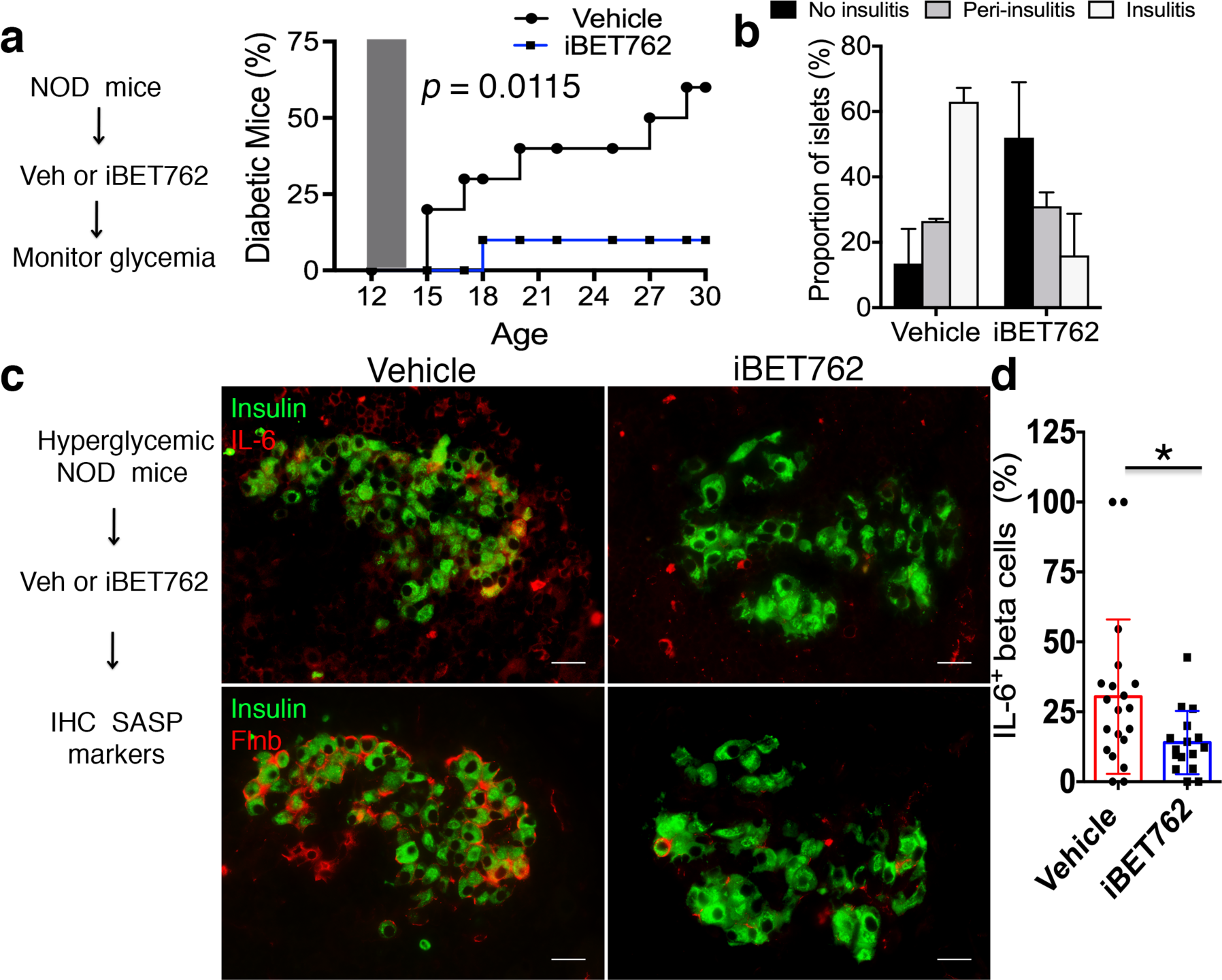
iBET-762 prevents diabetes and attenuates beta cell SASP in vivo. (a) 12-week NOD mice (n =10 per group) were administered vehicle or iBET-762 i.p. for 2 weeks and glycemia was monitored up to 30 weeks of age. Diabetes incidence curves shows frequency of hyperglycemic mice at each time point. Gray shaded area shows treatment window (12-14 weeks of age). *p* = 0.0115, Log-rank test. (b) Insulits grading in n = 30 islets taken from two euglycemic vehicle control and two euglycemic iBET-762 mice after the prevention study in (a) at 30 weeks. Error bars are SD. (c) Recent onset hyperglycemic NOD mice were administered vehicle or iBET-762 i.p. for one week, and then IHC was performed for SASP markers. Representative IHC of SASP factors IL-6 or Flnb with Insulin in pancreas sections from the indicated mice after treatment. Scale bars = 20 µm. (d) Quantification of IL-6^+^/Ins^+^ cells per islet in the hyperglycemic NOD mice treated as in (a). Each dot represents an islet, where n = 20 islets for a total of 735 beta cells were scored from three vehicle-treated mice, and n = 16 islets for a total of 891 beta cells were scored from three iBET-762-treated mice. *p < 0.05, two-tailed T-test.

We hypothesized that the protective effect of BET protein inhibition resulted at least in part from its inhibition of SASP in beta cells. Consistent with this, treatment of recent onset hyperglycemic NOD mice with iBET-151 also had a beneficial effect on beta cell function [21], which would be expected if this treatment affects the SASP program. To test whether BET inhibition diminished the beta cell SASP in vivo, recent onset hyperglycemic NOD mice were injected every day for one week with either vehicle or iBET-762 and pancreas sections were immunostained for SASP markers (Fig. 4c). Vehicle-injected hyperglycemic NOD mice showed a robust expression of IL-6 and Flnb in beta cells (Fig. 4c) consistent with beta cell SASP [3]. In contrast, islets from iBET-762-injected hyperglycemic mice showed fewer beta cells expressing IL-6 or Flnb (Fig. 4c). The frequency of IL-6^+^ beta cells per islet was variable in hyperglycemic mice, similar to our previous observations of SASP heterogeneity in islets of euglycemic NOD mice [3]. Nevertheless, quantifications showed an approximately 50% reduction in the average frequency of IL-6^+^ beta cells per islet in iBET-762-treated hyperglycemic mice relative to vehicle controls (30% in vehicle versus 14% in iBET-762, Fig. 4d). Taken together, these results demonstrate that BET inhibition with iBET-762, which is sufficient to prevent diabetes in NOD mice also blunts SASP in beta cells in vivo.

## 4. Discussion

In summary, our findings implicate BET proteins as crucial transcriptional activators of SASP in the beta cells of mice and humans. Our findings also indicate that in NOD mice the protection afforded by BET inhibitors against diabetes involves attenuation of SASP in beta cells. However, it is important to recognize that since BET proteins are also expressed in some immune cell types, where they activate genes for immune responses, BET inhibitors can also block immune inflammatory responses unrelated to SASP [20]. In addition, like their predecessor JQ1, second-generation BET inhibitors such as iBET-151 and the more potent derivative iBET-762 affect the chromatin binding of all of the paralogues [22] and there may be different contributions of BRD2, BRD3 and BRD4 to SASP activation in beta cells versus inflammatory responses in immune cells. Regardless, the net-effect of BET inhibition in the NOD model is remarkably beneficial perhaps as a result of both diminishing SASP in beta cells and reducing inflammatory gene expression in immune cells [21].

Transcriptional inhibition of beta cell SASP genes with iBET-762 had a corresponding downstream effect on SASP factor secretion and paracrine activity. Remarkably, BET inhibition also affected the extent of persistent DDR and SASP in human islets exposed to DNA damage, revealing a conservation of this mechanism in human islets. Inhibition of BRD4 in human fibroblasts after the induction of senescence also inhibits activation of SASP [8] consistent with our findings in human islets. While we have focused the current study on BRD4, genetic approaches will be required to delineate contributions of BRD2 and BRD3 to SASP activation in mouse and human beta cells.

The mechanism by which BET proteins are recruited to SASP gene regulatory regions in beta cells remains to be determined. BRD4 interacts with NF-kB to facilitate transcriptional activation of SASP genes in fibroblasts [8,23]. Although this pathway may also operate in during SASP in beta cells, there may also be beta cell-specific transcription factors induced during senescence that recruit BET protein activity. SASP beta cell-specific transcription factors that interact with BRD4 would be attractive targets for designing small molecules to inhibit SASP gene expression specifically in beta cells. Unlike using BET inhibitors, this approach would disentangle the effects of SASP inhibition in beta cells from the inhibitory effects on inflammatory responses in immune cells. Selective targeting of transcriptional and epigenetic regulators such as the BET protein family using small molecule probes is emerging as a powerful approach to treat a variety of diseases resulting from their aberrant activity [22]. As BRD4 is also expressed in human beta cells and contributes to the SASP transcriptional program in human islets, our work suggests that this concept may also be relevant to the prevention and/or treatment of T1D in humans. In conclusion, targeted inhibition of the SASP transcriptional program may provide a new opportunity to develop therapies for T1D.

## Acknowledgements

We would like to thank Mark Atkinson for assistance with the nPOD repository. P.J.T. was supported by a grant from the Diabetes Research Connection and a postdoctoral fellowship from the Hillblom Foundation. This work was supported by start-up funds from the UCSF Diabetes Center to A.B.

## Author Contributions

P.J.T., A.S. and A.B. designed the study. P.J.T., A.S. and C.A. carried out the experiments. P.J.T. and A.B. wrote the manuscript.

